# Behavioral dysregulation and monoaminergic deficits precede memory impairments in human tau-overexpressing (htau) mice

**DOI:** 10.1101/2022.04.03.486898

**Authors:** Kanza M. Khan, Govindhasamy Pushpavathi Selvakumar, Nagalakshmi Balasubramanian, Ruixiang Wang, Samantha Pierson, Marco Hefti, Catherine A Marcinkiewcz

## Abstract

Alzheimer’s disease (AD) poses an ever-increasing public health concern as the population ages, affecting more than 6 million Americans. AD patients present with mood and sleep changes in the prodromal stages that may be partly driven by loss of monoaminergic neurons in brainstem, but a causal relationship has not been firmly established. The goal of the present study was to evaluate depressive and anxiety-like behaviors in a mouse model of human tauopathy (htau mice) at 4 and 6 months of age prior to the onset of cognitive impairments and correlate these behavior changes with tau pathology, neuroinflammation, and monoaminergic dysregulation in the DRN and LC. We observed depressive-like behaviors at 4 months of age in male and female htau mice and hyperlocomotion in male htau mice. At 6 months, male htau mice developed anxiety-like behavior in the EZM, whereas hyperlocomotion had resolved by this time point. Depressive-like behaviors in the social interaction test persisted at 6 months but were resolved in the sucrose preference test. There was also a significant reduction in number and density of 5-HT-immunoreactive neurons in the rostral DRN in htau mice at 4 months and 5-HT neuronal density was negatively correlated with the intensity of phosphorylated tau staining in this subregion. Additionally, we found evidence of microglial activation in the mid and caudal DRN and astrocytic activation in the rostral DRN. 5-HT neuronal activity was reduced in the DRN and accompanied by downregulation of *Tph2* and *Sert*, whereas genes that promote neuroinflammation and tau phosphorylation were upregulated. Finally, there was enhanced ptau202/205 staining and microglial activity in the LC of htau mice and reduced TH optical density, although the number and density of TH+ neurons were not altered. In total, these results suggest that tau pathology in the DRN and the resulting loss of serotonergic neurotransmission may drive depressive-like behaviors in the early stages of AD, whereas anxiety-like behaviors develop later and may result from neurodegeneration in other regions.

## INTRODUCTION

Alzheimer’s disease (AD) is a devastating age-related neurodegenerative disease that afflicts a large proportion of individuals aged 65 and older[1]. Recent evidence suggests that neurofibrillary tangles (NFT) and amyloid plaques (AP) may develop in brainstem nuclei including the dorsal raphe nucleus (DRN) and locus coeruleus (LC) before the hippocampus and cortex, leading to behavioral changes in the prodromal stages of AD [2–8]. The DRN contains a large population of serotonin (5-HT) neurons that project to the forebrain and regulate mood, sleep and reward-seeking behaviors, all of which are perturbed in AD [9–12]. Although the role of the DRN in depressive-like behaviors has been extensively reported in the literature[10,13–15], a causal link between DRN tauopathy and depression in AD has not been rigorously established. The LC is also associated with depression and anxiety-like behaviors in rodent models[16,17], with studies suggesting that even a minimal loss of noradrenergic neurons can lead to depressive behavior[18]. The onset of tau pathology in these brainstem nuclei in AD is also accompanied by depletion of 5-HT and noradrenergic (NA) neurons [19–23], suggesting that monoaminergic depletion may be driving the early non-cognitive symptoms of AD.

The present study utilizes a mouse model (htau) that recapitulates human tauopathy to define the underlying neural substrates associated with early behavioral changes in AD. The htau mouse model overexpresses all six isoforms of the human microtubule-associated protein tau (*MAPT*) gene under human regulatory elements, so tau pathology develops in a more naturalistic fashion rather than being driven by an exogenous promoter. These htau mice were previously found to exhibit widespread tau pathology and cognitive deficits around 12 months of age [24,25], but non-cognitive symptoms such as anxiety and depressive-like behaviors have not been extensively studied. We examined depressive and anxiety-like behaviors in htau mice at 4 and 6 months of age using a battery of tests including the elevated plus maze (EPM) and elevated zero maze (EZM), open field, social interaction test, sucrose preference test, forced swim test. Spatial memory was assessed in the Barnes maze at 6 months of age. We also examined monoaminergic markers (5-HT, TH), tau phosphorylation (AT8), and markers of microglia and astrocytic activity (Iba-1, GFAP) in the DRN and LC with immunohistochemistry. RT-qPCR was used to identify potential biomarkers of brainstem neuropathology with a panel of genes involved in monoamine biosynthesis and signaling, neuroinflammation, and protein aggregation. Finally, we examined 5-HT neuronal activity in brain slices using patch clamp electrophysiology as a proxy for 5-HT neuronal function and output. Overall, our results suggest that htau mice exhibit depressive behaviors at 4 months that coincide with deficits in 5-HT neuronal activity, reduced 5-HT levels, and increased neuroinflammation and tau accumulation in the DRN. Anxiety-like behaviors develop later at 6 months and may be caused by neurodegeneration of TH neurons in the LC or other brain regions.

## MATERIALS AND METHODS

### Animals

Male and female C57BL/6J mice (Jackson Labs #000664) and htau +/- mice (Jackson Labs #005491) containing a transgene that encodes the human *MAPT* gene were used in this experiment. Htau -/- (global tau knockout) mice were used as negative controls in RT-PCR and Western blot validation of tau splice isoforms. Mice were housed in a temperature-and humidity-controlled, AALAC-approved vivarium at the University of Iowa with *ad libitum* access to food and water.

#### RT-PCR quantification of tau splice isoforms

##### RNA extraction

Total RNA was extracted from cortical tissues of WT, htau +/- and htau -/- mice brains using TRIzol reagent (Ambion, Life Technologies, USA) as described previously [26]. The DNA contaminants from the extracted RNA were eliminated using a DNA-free™ DNA Removal Kit (Life Technologies, USA). The purity and quantity of RNA were checked using NanoDrop 1000 spectrophotometer (Thermo Fisher Scientific, USA) for downstream experiments.

##### cDNA synthesis and RT-PCR analysis

To examine the 3R (3 repeats; 293bp) and 4R (4 repeats; 390bp) *MAPT* isoform, we performed RT-PCR using specific primers reported previously [27,28]. The total RNA extracted was reverse transcribed using the iScript™ cDNA Synthesis kit (Bio-Rad Laboratories, CA, USA) with a thermal profile: 25 °C for 5 min, 45 °C for 20 min, and 95 °C for 1 min. Following this, RT-PCR was performed using the thermal profile: 95 °C for 3 min followed by 40 cycles of 95 °C for 30 s, 60 °C for 30 s, and 72 °C for 1 min and final extension at 72 °C for 5 mins. After RT-PCR, the products were resolved in a 2% agarose gel and imaged under UVPTM gel imaging system (UVP LLC, CA, USA).

#### RT-qPCR analysis of monoaminergic, inflammatory and tau phosphorylation genes

Anatomically distinct brain regions including the DRN and LC tissues were micro-punched from WT and htau+/- carrier brains and total RNA was isolated. The concentration and purity was checked using a NanoDrop 1000 and reverse transcribed to cDNA. RT-qPCR for the target genes was performed using SYBR green qPCR master mix (Bio-Rad Laboratories, USA) and specified primers (Table 1) on a CFX96™ Real-time-PCR System (Bio-Rad Laboratories, USA). The thermal profile used for RT-qPCR was 95 °C for 10 min, followed by 40 cycles of 95 °C for 30 s, 60 °C for 30 s, followed with a melt curve analysis profile (60 °C to 95 °C in 0.5 °C increments at a rate of 5 s/step). Fold changes in the mRNA levels were determined for each gene after normalizing with β-actin Ct values using the fold change 2^−ΔΔCT^ method [29]. Results are represented as fold changes in the mRNA levels (± SEM).

### Western blots analysis of tau isoforms

For detection of total human tau, tissues were weighed and homogenized in RIPA buffer at 150 μl/mg tissue using a sonicator. For detection 3R and 4R isoforms, tissues were homogenized in high-salt (HS) lysis buffer using a bead mill homogenizer [24,30–34]. The homogenate was then centrifuged at 130,000 *x g* for 30 minutes at 4□C. The supernatant was removed, and the tissue pellet was resuspended in 1% Triton X-100 in HS buffer and centrifuged at 130,000 *x g* for 30 minutes at 4□C. The supernatant was removed, and this process repeated a second time. To isolate the soluble and insoluble fraction, the pellet was resuspended in 1% of Sarkosyl was in HS buffer and incubated for 30 minutes in a water bath at 37□C. This suspension was then centrifuged at 130,000 *x g* for 30 mins at 4°C. The supernatant was removed (Sarkosyl soluble fraction), and the pellet re-suspended in urea buffer (0.5 ml of 4 M urea, 2% SDS, and 25 mM Tris-HCl, pH 7.6; Sarkosyl insoluble fraction). Protein concentration was measured using the Pierce™ 660nm Protein Assay Kit (ThermoFisher Scientifc) following the manufacturer’s protocol.

Extracted protein lysate samples were mixed with Laemmli buffer (Bio-Rad Laboratories, USA) and denatured at 95°C for 5 mins. Samples were then loaded into a precast 10-well 10% TGX SDS-PAGE gel (BioRad Laboratories, USA), and proteins were transferred to a PVDF membrane by using semi-dry Trans-Blot Turbo Transfer System (BioRad Laboratories, USA). Then the membranes were transferred into blocking solution (10% BSA or skimmed milk powder in PBS with 0.1% Tween20). Membranes were probed with primary antibodies as follows: HT7 (Total human tau; 1:3000), RD3 (3R; 1:200), and RD4 (4R; 1:200). HT7 was obtained from ThermoFisher and RD3 and RD4 were generous gifts from Gloria Lee. The following day, all membranes were probed with appropriate secondary antibodies (IRDye 680RD; 1:2000 or IRDye 800CW; 1: 3000; Li-Cor Biotechnology, USA) for 60 mins followed by 3 washes with PBS-T. Membranes were then imaged with a Sapphire™ Biomolecular Imager (Azure Biosystems, Inc).

### Immunohistochemistry (IHC)

Mice were anesthetized with Avertin and transcardially perfused with 30 ml 0.01 M PBS followed by 30 ml 4% paraformaldehyde (PFA). Brains were extracted, fixed in PFA for 24 h at 4°C, and stored in 0.02% Sodium Azide/PBS at 4°C. After the gradient sucrose treatment (10, 20 and 30%) brain tissues were fixed in a cryomold with O.C.T. fixative then moved to −80 for overnight. Then next day cryomolds were transferred into −20 □C for 4 hr and then sectioned at 25 μm using a Leica cryostat maintained at −15 to −20 □C. Slices were stored at 4°C in a cryoprotectant solution (3% sucrose, 0.10% Polyvinyl-pyrrolidone (PVP-40), 0.05% of 0.1M Phosphate Buffer, and 0.03% ethylene glycol). For each region of interest (ROI), 3-4 slices were used across the caudal-rostral axis. Slices were washed in PBS and incubated in 0.5% Triton X-100/PBS for 30 min, blocked in 10% Normal Donkey Serum in 0.1% Triton X-100/PBS and then incubated with the respective primary and secondary antibodies (Table 2). When staining for AT8, Mouse on Mouse blocking buffer (3% final volume; Vector Laboratories) was added to the blocking solution to reduce non-specific binding to endogenous mouse IgG. Slices were subsequently washed in PBS, mounted on glass slides and coverslipped with Vectashield mounting media (Vector Laboratories, Inc.).

Confocal z-stacks (1 μm) were captured on an Olympus FV3000 laser scanning confocal microscope (20 sections/z-stack) and converted to maximum projection images using Image J software. Images were analyzed by trained researchers blind to experimental conditions to obtain cell counts per unit area, percent immunoreactive area, and optical density using ImageJ. The optical density was estimated by first converting images to an 8-bit gray scale image, and performing a background correction. Optical density calibration was performed with a 21-step tablet (available from ImageJ) using the Rodbard function. Following optical density calibration, mean gray scale values were recorded from the ROIs with an effort to avoid obvious artifacts. Percent immunoreactive area was performed on the ROIs of thresholded images. For each image, the ROI was drawn according to the shape of the region based on the atlas and the monoaminergic signal. ROIs were drawn with the aid of the 5-HT or TH stain, and then applied to AT8, iba-1, or GFAP channels.

### Slice Electrophysiology

#### Brain Slice Preparation

Deeply anesthetized mice were intracardially perfused with ice-cold, oxygenated modified artificial cerebrospinal fluid (aCSF) containing the following (in mM): 110 choline-Cl, 2.5 KCl, 7 MgSO_4_, 0.5 CaCl_2_, 1.25 NaH_2_PO_4_, 26.2 NaHCO_3_, 25 glucose, 11.6 Na-ascorbate, 2 thiourea, and 3.1 Na-pyruvate (pH: 7.3 – 7.4; osmolality: 300 – 310 mOsmol/kg). The brains were quickly dissected and coronal midbrain slices containing the DRN (300 μm) were obtained using a vibratome (VT1200S; Leica Biosystems, Wetzlar, Germany). The brain slices recovered at 34 °C for 30 min in a chamber containing the choline-Cl-based aCSF described above, continuously bubbled with 95% O_2_/5% CO_2_. After the initial recovery, brain slices were transferred to a different modified aCSF at room temperature saturated with 95% O_2_/5% CO_2_ for at least 1 hour before recordings started. The holding aCSF contained the following (in mM): 92 NaCl, 2.5 KCl, 2 MgSO_4_, 2 CaCl_2_, 1.25 NaH_2_PO_4_, 30 NaHCO_3_, 20 HEPES, 25 glucose, 5 Na-ascorbate, 2 thiourea, and 3 Na-pyruvate (pH: 7.3 – 7.4; osmolality: 300 – 310 mOsmol/kg).

#### Electrophysiological recordings

During recordings, slices were continuously perfused (2 ml/min) with standard aCSF containing the following (in mM): 124 NaCl, 4 KCl, 1.2 MgSO_4_, 2 CaCl_2_, 1 NaH_2_PO_4_, 26 NaHCO_3_, and 11 glucose (osmolality: 300 – 310 mOsmol/kg), saturated with 95% O_2_/5% CO_2_ and maintained at 30 ± 1°C. Patch electrodes (3 – 5 MΩ) were filled with a solution containing the following (in mM): 135 K-gluconate, 5 NaCl, 2 MgCl_2_, 10 HEPES, 0.6 EGTA, 4 Na_2_-ATP, and 0.4 Na_2_-GTP (pH: 7.35; osmolality: 288 – 292 mOsmol/kg). In order to identify 5-HT neurons *post hoc* by IHC, biocytin (2 mg/ml; Tocris Bioscience, Bristol, UK) was added into the internal solution. Only Tph2-positive neurons shown by IHC were included in data analysis. Neurons were visualized via an upright microscope (BX51W1; Olympus, Tokyo, Japan) accompanied by an infrared differential interference contrast imaging system. Membrane currents were amplified with a Multiclamp 700 B amplifier (Molecular Devices, San Jose, CA, USA), filtered at 3 kHz, and sampled at 20 kHz with a Digidata 1550B digitizer (Molecular Devices). Data were acquired via the pClamp 11 software (Molecular Devices). Access resistance was monitored online and changes greater than 10% would lead to exclusion of the recordings from data analysis.

To examine neuronal intrinsic excitability, recordings were conducted in the current clamp mode. In some experiments, currents were injected into cells to hold the membrane potential at −70 mV. Input resistance was assessed by the change in membrane potential when the cell was hyperpolarized with −100 pA current injection. Rheobase reflected the minimal currents needed to evoke action potentials. Numbers of evoked action potentials (i.e., spike numbers) were recorded after injecting positive currents for 250 ms at 10 pA incremental steps (0 – 200 pA).

### Behavior

Anxiety- and depressive-like behaviors in C57BL/6J and htau +/- mice were evaluated at 4 and 6 months of age. All behavior tests were performed between 9am-3pm and were recorded with an overhead or side-view camera integrated with Media Recorder or Ethovision video tracking software (Noldus Information Tech, Inc.). Videos were scored by an experimenter blinded to animal genotypes. Unless otherwise noted, all behavior tests were scored using Ethovision.

#### Elevated plus maze

At 4 months of age, animals were tested in the EPM to evaluate anxiety-like behaviors[35]. Briefly, the maze was 60cm above the floor and consisted of two ‘open’ arms, two ‘closed’ arms (5 × 35 cm), and a neutral starting zone (5 × 5 cm). Overhead LEDs were used to maintain lux in the open arms at 20 lux and <5 lux in the closed arms. The closed arms had tall dark walls that allowed animals to hide. At the beginning of the test, animals were placed in the neutral zone and allowed to freely explore the maze for 5 min. Distance traveled, and time spent in the open arms vs closed arms were calculated using Ethovision XT14.

#### Elevated zero maze

At 6 months of age, animals were tested in the EZM to evaluate anxiety-like behaviors in a novel arena [36–38]. Briefly, the circular maze (outer diameter: 52cm) was elevated 60cm above the ground and consisted of two alternating open and closed corridors (width: 5cm). Overhead LEDs maintained the lux in the open corridors at 20 lux and <5 lux in the closed corridors. At the beginning of the test, animals were placed in the open corridor, facing the closed corridor and allowed to freely explore the arena for 5 min. Behavior was recorded by an overhead camera. Distance traveled, and time spent in the open arms vs closed arms were calculated using Ethovision XT14. The open arm preference, and probability of entering the open arms were calculated to determine the anxiety-like behavior.

#### Open field test

Mice were placed in the corner of a 50 × 50 × 25cm plexiglass arena and allowed to freely explore the arena for 30min. The open field test was performed at 4- and 6-months of age to evaluate locomotor and exploratory behavior. The total distance traveled (cm), time spent in the center of the arena, and time spent in the corners of the arena were measured through Ethovision XT14. The center of the open field was defined as the central 15% of the arena.

#### Social interaction test

The social interaction test was performed in 4- and 6-month-old mice, and was performed as previously described [39,40]. Briefly, the animal was placed in the central chamber of a 3-chamber arena (20 lux) and allowed to explore the environment for 10 min. Following this, a novel C57BL/6J mouse of the same sex as the test mouse was placed in one of the side chambers under a metal cage; an empty metal cage was placed in the alternate side chamber. The test mouse was allowed to explore the environment and interact with the stranger mouse for 10 min. The location of the stranger mouse was alternated between the right- and left-chamber to control for any potential side preferences. The total time spent interacting with the stranger mouse, and with the novel object (empty cage) was scored by an experimenter blinded to animal genotypes.

#### Sucrose preference test

The SPT was performed over four days and used to evaluate the degree of anhedonia of htau mice at 4 and 6 months of age [41,42]. Animals were transferred to PhenoTyper observation cages (Noldus) that were fitted with two sipper bottles and Lickometers (to measure the number of licks the animal makes to a bottle). Animals had ad libitum access to tap water or a 5% sucrose solution (ThermoFisher) for 1-hour. The placement of sucrose and water bottles were alternated each day to control for any side preference. The number of approaches to the water bottle and the sucrose bottle was measured through Ethovision XT14.

#### Barnes maze

Spatial learning and cognitive deficits were evaluated in 6-month-old C57 and htau mice as described previously [43–45]. The Barnes maze was a 5-day protocol[46], with environment habituation on day 1, training sessions on days 2 and 3, rest on day 4, and a probe trial on day 5. The maze was a gray circular arena (diameter: 91cm), consisting of 20 equally divided holes, and was elevated 93 cm above the ground. The room was well lit and visual cues were present on the walls.

On Day 1, animals were guided to the predetermined ‘goal box’, which had been fitted with an escape chamber. In contrast, the remaining 19 (non-target holes) were not fitted with any chambers, and animals could see the ground below.

Over days 2 and 3, animals underwent a total of five operant conditioning training trials. A loud buzzer noise (~100 dB) was played while animals explored the environment. Once the animal found the goal box and entered the escape chamber, the buzzer noise was turned off and animals were allowed to rest for 1-min before being returned to a holding cage. If an animal did not find the goal box/escape chamber within a 2-min trial, they were guided to the escape chamber as they had been on Day 1. Inter-trial interval was 30 min. Animals had 3 training trials on day 2, and 2 trials on day 3.

On the probe day, the escape chamber was removed from the goal box. Animals were placed on the maze platform, and the conditioned stimulus (buzzer sound) was presented. Animals were allowed to explore the environment for 2 min, and behavior was recorded by an overhead camera.

Recordings were scored through Ethovision XT14 by a researcher blind to animal genotypes. The number of visits to the goal box & non-target holes were measured along with the latency to approach the goal box, and time spent in the target quadrant.

#### Forced Swim Test

Depressive like behavior was evaluated at 4- and 6-months in the forced swim test. Mice were gently placed in a tall cylinder filled with 24-25°C tap water (32 cm height x 20 cm diameter, water height: 25cm) for 6 minutes. After the test, animals were placed in a clean cage under a heating lamp for 5-minutes to warm them and allow them to dry off. Animal behavior was recorded from a side-view camera and analyzed with Ethovision XT14. The 6-min videos were divided into two bouts: a pretest (1-2min) and a test (3-6) phase. The latency to the first immobile bout, frequency of immobile bouts, and duration of each immobile bout was evaluated.

## RESULTS

### Htau mice express 3R and 4R isoforms of tau

In this study, we used htau +/- mice (Jackson Labs strain # 005491) that had been backcrossed to a mouse tau knockout line (Mapt<tm1(EGFP)Klt). These mice are on a C57BL/6J background, so C57BL/6J mice were used as controls throughout this study as in previous reports [24,25,47]. Htau mice express all six isoforms of human tau including the 3R and 4R isoforms and are thought to represent a more naturalistic model of AD with late-onset cognitive impairment[25]. We first confirmed with RT-PCR that our htau +/- mice expressed human tau isoforms with primers spanning exons 2 and 3 (three isoforms) and exons 9-11 (3R and 4R) of the human tau gene. C57BL/6J and htau -/- mice lacked these amplification products. On the other hand, primers spanning exons 2-3 of the mouse tau gene produced bands in C57BL/6J mice that were absent in htau +/- and htau -/- mice (Figure 1A). C57BL/6J mice contained only the mouse tau fragment while both human and mouse tau fragments were absent in htau -/- mice. In a separate cohort, we extracted total protein for immunoblotting analysis and probed for human tau in htau +/-, C57BL/6J, and htau -/- mice with an HT7 antibody and confirmed that human tau was only present in the htau +/- mice (Figure 1Bi). Next, we isolated Sarkosyl soluble and insoluble fractions from these mice and probed for 3R and 4R isoforms with specific antibodies (RD3 and RD4, respectively). Here we observed 3R isoforms in both Sarkosyl-soluble and insoluble fractions in htau +/- mice, but not in C57BL/6J or htau -/- mice (Figure 1Bii)

**Figure 1:**
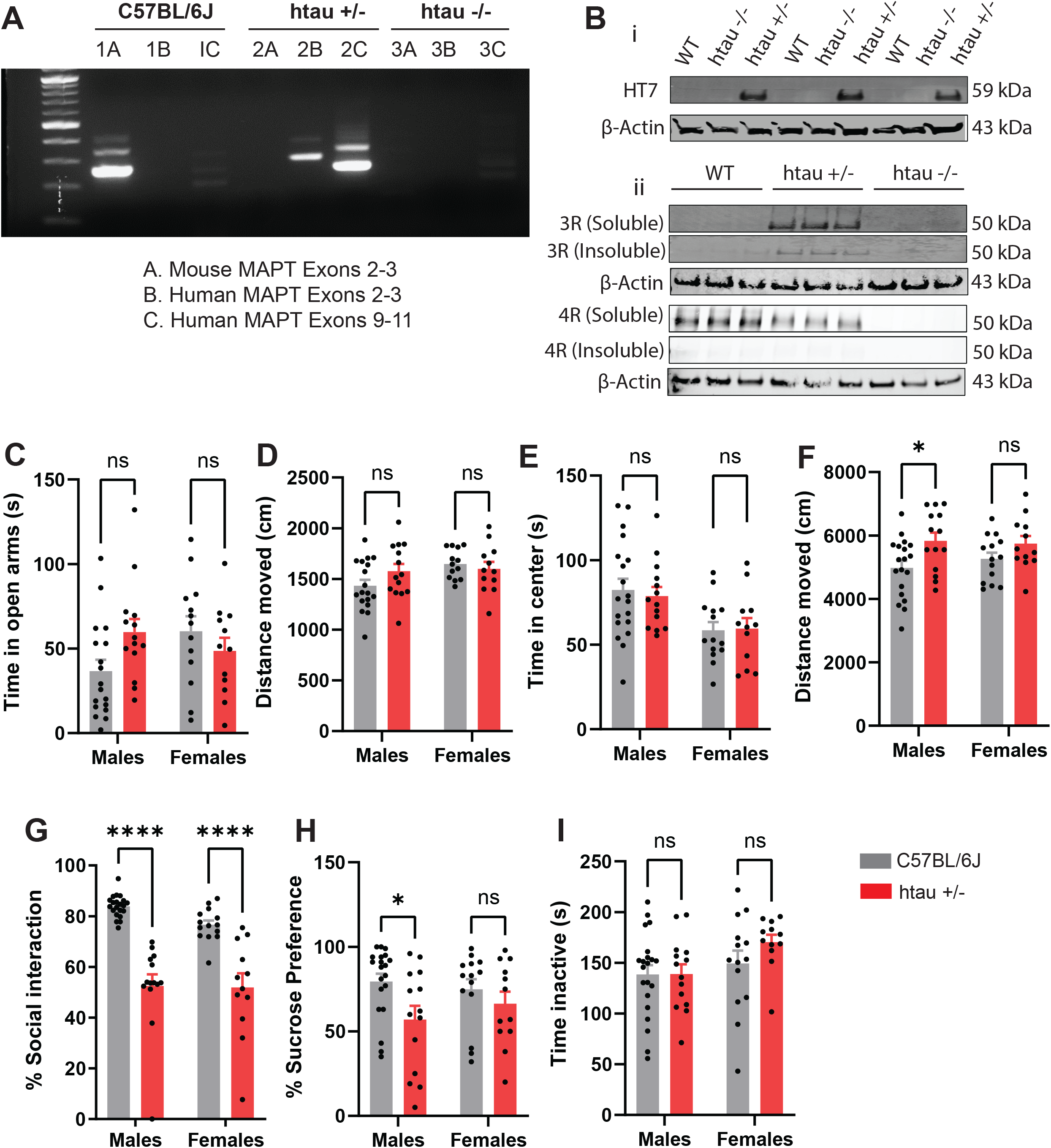
Htau mice exhibit depressive-like behaviors at 4 months of age. (A) Mouse and human tau splice isoforms in C57BL/6J, htau +/- and htau -/- mice. The human-specific primers only amplified products in htau +/- while mouse-specific primers only amplified products in C57BL/6J mice. (B) Protein expression of human tau (HT7) in total protein extracts (i) and 3R and 4R tau isoforms in Sarkosyl soluble and insoluble fractions from C57, htau +/- and htau -/- mice (ii). Only htau +/- mice contained the 3R tau isoform in detergent insoluble fractions and human tau. Both htau +/- and C57 mice contained the 4R isoform in soluble fractions. (C-D) Behavior in the EPM, (E-F) open field, (G) social interaction test, (H) sucrose preference test and (I) forced swim test in 4-month-old male and female C57 and htau +/- mice. These results indicate depressive-like behaviors in htau mice relative to C57 mice. *p<0.05, ****p<0.0001.

### Htau mice exhibit depressive-like behaviors by 4 months of age and anxiety-like behaviors by 6 months

We next asked whether htau mice exhibit behavioral phenotypes reminiscent of prodromal AD by 4 months of age, which coincides with the appearance of hyperphosphorylated tau in the DRN but prior to the onset of cognitive impairments [25,47]. Male and female htau and C57BL/6J mice were tested in the EPM, open field, social interaction test, sucrose preference test, and forced swim test at this time point. We found a significant interaction between sex and genotype in time spent in the open arms of the EPM at 4 months (F_1,53_=4.92, p<0.05), but post-hoc comparisons between htau and C57BL/6J mice were non-significant in both males and females. No group differences in locomotor activity were observed in this assay (Figure 1C-D). There was a main effect of sex in time spent in the center of the open field (F_1,55_=12.25, p<0.001) but no sex x genotype interaction, suggesting that female mice generally exhibited more anxiety-like behavior in this test. Htau mice also exhibited hyperlocomotion in the open field (Main effect genotype: F1,55=8.32, p<0.01) although Bonferroni post-tests were only significant in males (t_55_=2.77, p<0.05) (Figure 1E-F). Interestingly, both sexes exhibited significant depressive-like behaviors in the social interaction test (Main effect of genotype: F_1,57_=74.37, p<0.001; Bonferroni post-tests t_57_=7.32, p<0.01 for males and t_57_=5.06, p<0.001 for females) (Figure 1G), whereas sucrose preference was only reduced in male mice (Main effect genotype (F_1,56_=5.89, p<0.05; Bonferroni post-test: t_56_=2.65, p<0.05) (Figure 1H). Overall, there was no effect of genotype in the forced swim test, but female mice spent more time immobile than males (Main effect of sex: F_1,57_ = 4.22, p<0.05) (Figure 1I).

We also examined anxiety and depressive-like behaviors in these mice 6 months of age, except that the EZM was substituted for the EPM. Here we found a significant effect of genotype in time spent in the open area of the EZM (F_1,56_=12.21, p<0.001), with male htau mice exhibiting a significant reduction in open area time (Bonferroni post-tests: t_56_=3.11, p<0.01) (Figure 2A). Locomotor activity did not significantly differ between groups in this test (Figure 2B). There was also a significant effect of genotype in time spent in the center of the open field (F_1,55_=5.15, p<0.05) that is suggestive of enhanced anxiety-like behavior, although post-hoc comparisons were non-significant in both sexes (Figure 2C). Locomotor activity in the open field did not differ significantly between groups, which contrasted with the hyperlocomotive phenotype observed at 4 months (Figure 2D). Depressive-like behaviors in the social interaction test persisted at 6 months in both sexes (Main effect of genotype: F_1,50_=94.73, p<0.001; Bonferroni post-tests: t_50_=8.23 for males and t_50_=5.70 for females) (Figure 2E), although the decrease in the sucrose preference noted at 4 months had resolved by this time point (Figure 2F). Similar to what we observed at 4 months, there was no effect of genotype in the forced swim test at 6 months, although female mice continued to spend more time inactive than males (Main effect of sex: F_1,57_=5.76, p<0.05) (Figure 2G). There was also no effect of sex or genotype on spatial memory in the Barnes Maze at 6 months (Figure 2H), which is consistent with previous reports that memory impairments in htau mice only become apparent at 12 months of age[25].

**Figure 2:**
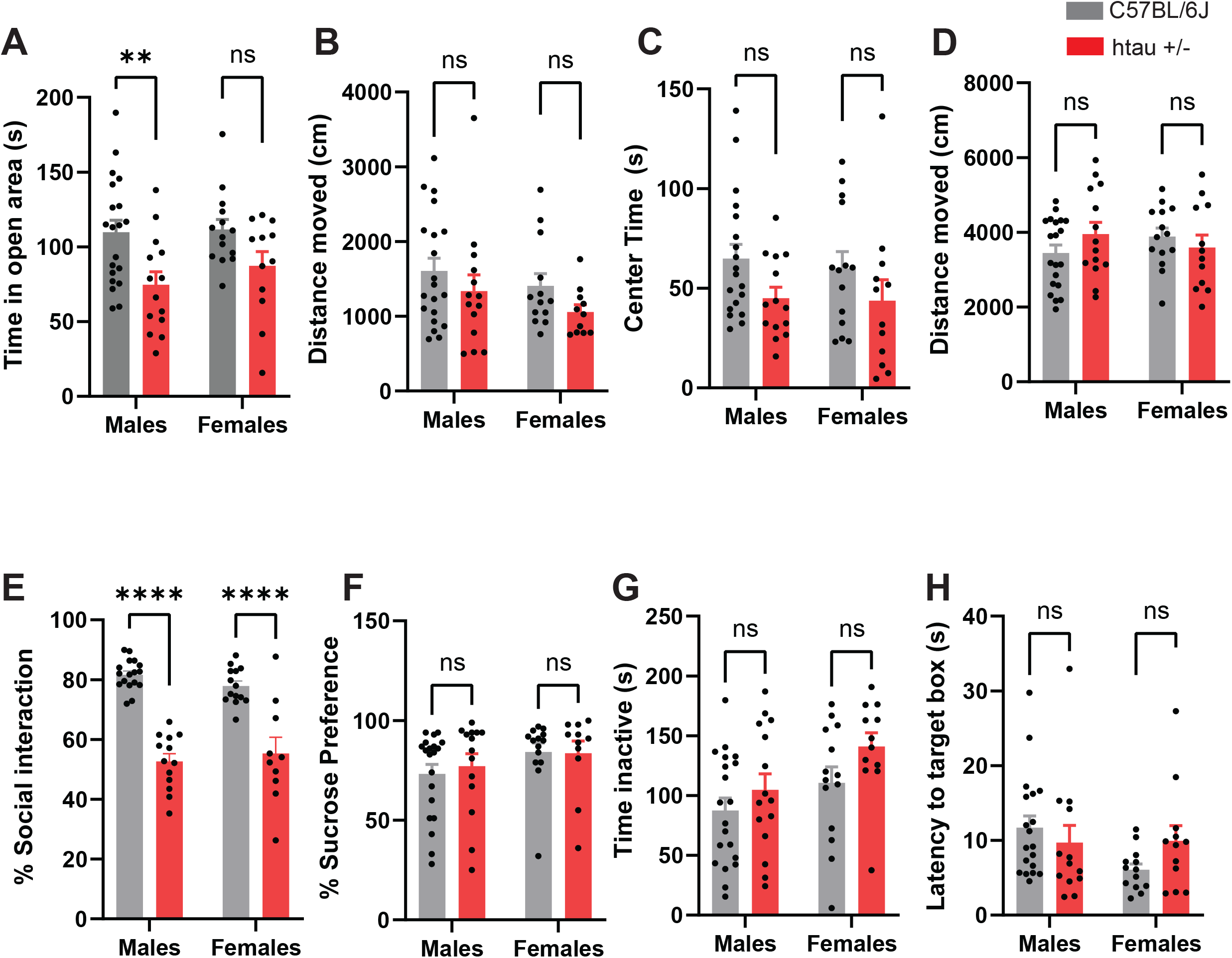
Htau mice exhibit anxiety and depressive-like behaviors at 6 months of age. (A-B) Behavior in the EZM, (C-D) open field, (E) social interaction test, (F) sucrose preference test, (G) forced swim test and (H) Barnes maze in 6-month-old male and female C57 and htau +/- mice. These results indicate anxiety-like behaviors in male htau mice at 6 months and a persistent social interaction deficit in both sexes. **p<0.01, ****p<0.0001.

### Monoaminergic depletion and hyperphosphorylated tau in the brainstem at 4 months

Next, we investigated whether loss of monoaminergic neurons in the DRN or LC at the 4-month mark might account for these behavioral phenotypes. Histological experiments were performed in separate cohorts of male htau and C57BL/6J mice to avoid any confounds of behavioral testing on monoamine levels. The DRN was subdivided into rostral, mid and caudal subregions which were previously found to have distinct forebrain projections and behavioral outputs[13,15,48,49]. Here we found a significant reduction in the density of 5-HT-immunoreactive (5-HT-IR) neurons (t_8_=3.60, p<0.01) and optical density of 5-HT (t_8_=3.40, p<0.01) in the rostral DRN, while in the mid and caudal DRN 5-HT neuronal and optical density was trending downward but not significantly altered (Figure 3A-D). We also used a ptau202/205 antibody (AT8) to assess tau pathology in the DRN and found there were trends toward increased AT8 immunoreactive area and optical density that did not reach significance in all subregions of the DRN (Figure 3E-F). However, we did observe a significant correlation between AT8 optical density and 5-HT neuronal density in the rostral DRN (R^2^= 0.5152, p<0.05), but not in other subregions (Figure 3G). Furthermore, there was colocalization between 5-HT neurons and AT8 in the DRN indicating the presence of tau inclusions in these neurons (Figure 3A).

**Figure 3:**
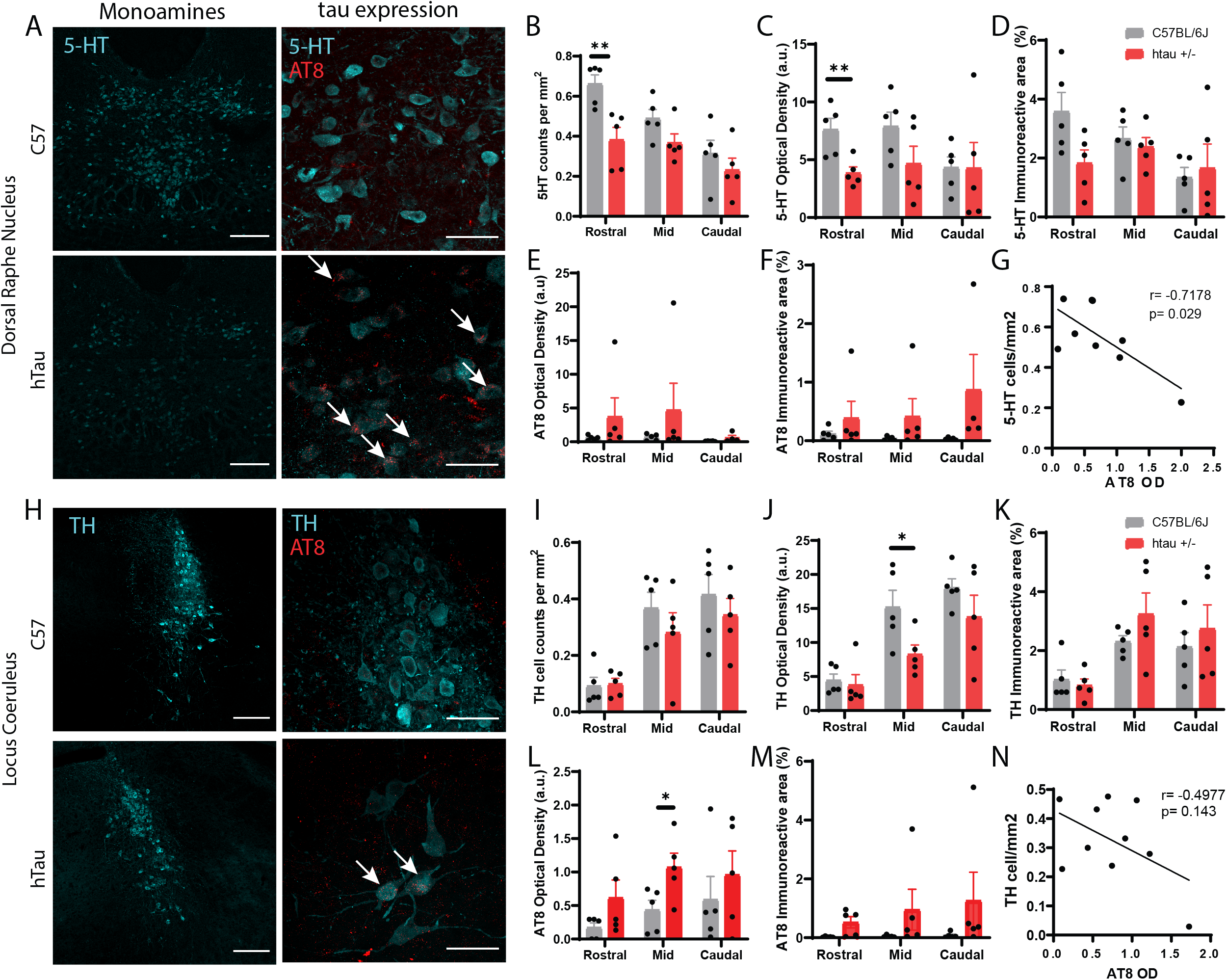
Reduction in 5-HT neuronal density correlates with an increase in phospho-tau in the DRN of htau mice at 4 months. (A) Representative confocal images of 5-HT immunostaining (20X; scale bar = 200 μm) and 5-HT/AT8 colocalization (60X; scale bar = 50 μm) in the DRN of C57 and htau +/- mice. (B) 5-HT cell counts/mm^2^, (C) optical density and (D) immunoreactive area in subregions of the DRN. (E) AT8 optical density, (F) immunoreactive area in subregions of the DRN, and (G) correlation analysis between 5-HT neuronal density and AT8 optical density in the rostral DRN. (H) Representative confocal images of TH immunostaining (20X; scale bar = 200 μm) and TH/AT8 colocalization (60X; scale bar = 50 μm) in the LC of C58 and htau +/- mice. (I) TH cell counts/mm^2^, (J) optical density, and (K) immunoreactive area in subregions of the LC. (L) AT8 optical density and (M) immunoreactive area in subregions of the LC and (N) correlation analysis between TH neuronal density and AT8 optical density in the mid LC. *p<0.05, **p<0.01. White arrows denote colocalization between AT8 and Tph2 or TH in the htau mice.

The LC was also divided into rostral, mid and caudal subregions for analysis. Surprisingly, there was no change in the neuronal density of TH-IR neurons in any subregion of the LC (Figure 3H-I). We did observe a reduction in the optical density of TH in the mid LC (t_8_=2.46, p<0.05), which is in line with clinical reports of reduced NE function in the prodromal stages of AD (Figure 3J-K). We also observed an increase in AT8 optical density in the mid LC (t_8_=2.51, p<0.05) which also has the highest number of TH-positive neurons (Figure 3L-M). There was no significant correlation between AT8 optical density and TH neuronal density or TH optical density in any subregion of the LC (Figure 3N).

It was surprising that we did not observe loss of TH-IR neurons despite the presence of hyperphosphorylated tau in this area and colocalization between phospho-tau and TH-positive neurons. By contrast, there was a decline in 5-HT-IR neuronal density in the DRN that correlated with AT8 immunoreactivity, suggesting that this neuronal population may be more susceptible to tau-induced neurodegeneration. On the other hand, we did not measure norepinephrine levels in the LC directly, and although there should be a positive correlation between tyrosine hydroxylase expression and norepinephrine synthesis, this may not always be the case.

### Glial cell activation in the DRN and LC of htau mice at 4 months of age

Clinical reports have suggested that neuroinflammation is a significant occurrence in AD and may play a role in the pathogenesis of the disease, promoting both tau aggregation and neurodegeneration[50,51]. Most studies focus on the role of microglia and astrocytes which are found in abundance at sites of amyloid and tau pathology, but their function in the etiology of AD in controversial. There is some evidence of a neuroprotective role, while others have found that glial activation can promote synaptic engulfment and release cytokines that can be toxic to neurons[52–55]. Mouse models of AD tend to support a neurodegenerative role of glia, and in the LC it was reported that loss of norepinephrine promotes inflammation while reducing microglia phagocytosis and clearance of β-amyloid plaques[56]. In the present study, we examined microglial and astrocytic markers Iba-1 and GFAP in the DRN and LC of 4-month-old male htau and C57 mice. The mid and caudal DRN displayed increased Iba-1 optical density (t_7_=2.74, p<0.05 and t_8_=2.76, p<0.05, respectively) (Figure 4A, 4C-E). The rostral DRN, which was the subregion with reduced 5-HT neuronal density, had a significant increase in GFAP optical density (t_8_=3.78, p<0.01) which is indicative of astrocytic activation (4B, 4F-H). There was also an increase in GFAP immunoreactive area in the caudal DRN (t_8_=2.85, p<0.05).

**Figure 4:**
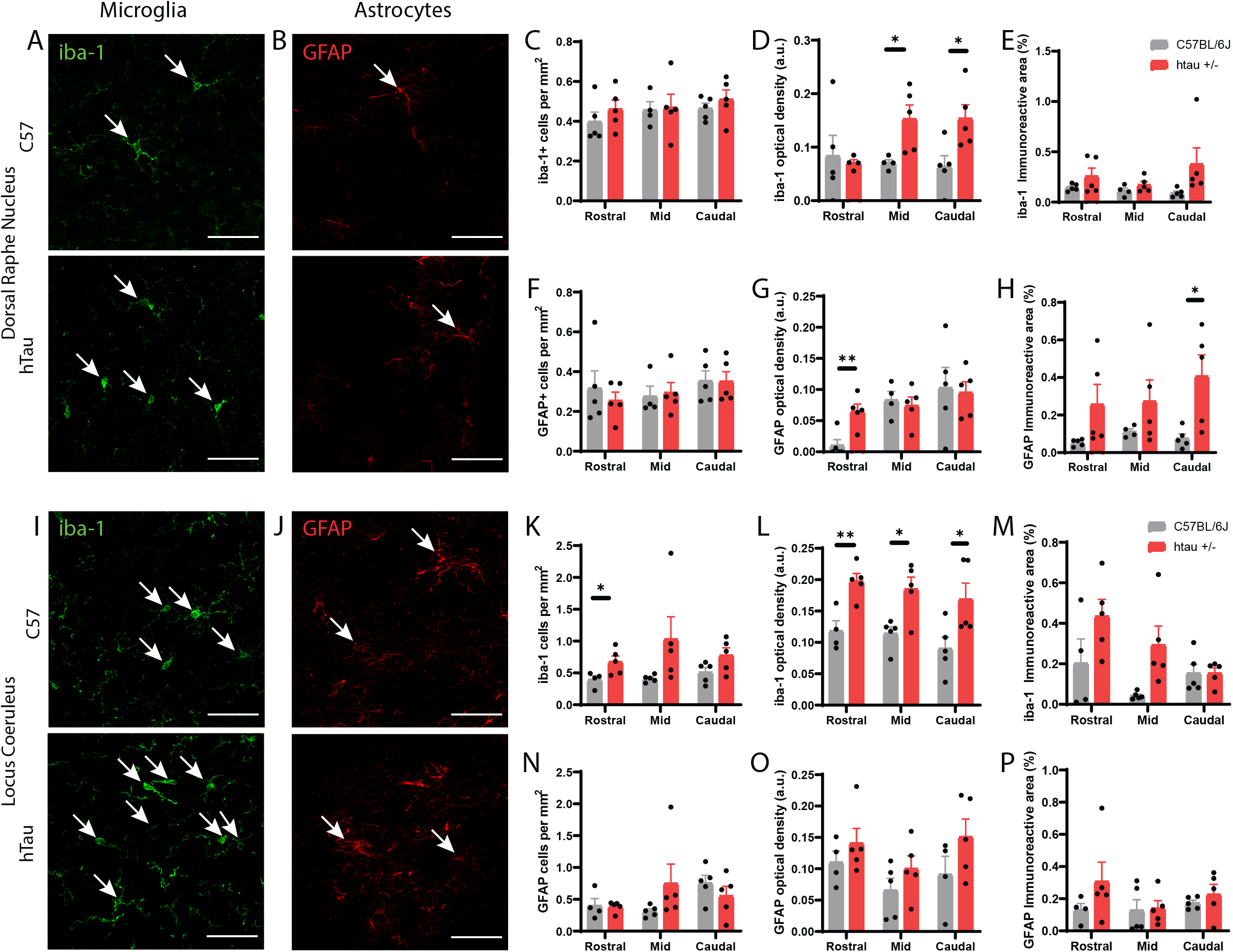
Glial activation in the DRN and LC in htau mice at 4 months. (A) Representative confocal images of Iba-1 and (B) GFAP immunostaining in the DRN of C57 and htau +/- mice (60X; scale bar = 50 μm). (C) Iba-1+ cell counts/mm^2^, (D) optical density, and (E) immunoreactive area in subregions of the DRN. (F) GFAP+ cells/mm^2^, (G) optical density, and (H) immunoreactive area in subregions of the DRN. (I) Representative confocal images of Iba-1 and (J) GFAP immunostaining in the LC of C57 and htau +/- mice. (K) Iba-1+ cells/mm2, (L) optical density, and (M) immunoreactive area in subregions of the LC. (N) GFAP+ cells/mm^2^, (O) optical density, and (P) immunoreactive area in subregions of the LC. *p<0.05, **p<0.01. White arrows denote Iba-1 +or GFAP+ cell bodies.

In the LC, there was a significant increase in Iba-1 optical density in all subregions of the LC (rostral: t_7_=4.02, p<0.01; mid: t_8_=3.26, p<0.05; caudal:t_8_=2.51, p<0.05) (Figure 4I, 4K-M). Additionally, the neuronal density of Iba-1+ microglia in the rostral LC was increased in htau mice (t_7_=2.43, p<0.05). In contrast to the DRN, there was no change in GFAP staining in any LC subregion indicating that astrocytes were not in an activated state (Figure 4J, 4N-O). The increase in Iba-1 in the LC is interesting in view of previous reports that microglia may migrate and become phagocytic in response to NE stimulation[56]. While we did not look at microglial migration or phagocytosis directly, it does suggest that this may be a response to the presence of tau pathology in this region. The intact TH+ neurons in the LC at this time point may enable NE release to increase tau clearance via microglial phagocytosis.

### Altered expression of genetic markers of monoaminergic transmission, neuroinflammation and protein aggregation in brainstem nuclei

The accumulation of tau pathology in the DRN and LC of htau mice may have downstream effects on monoaminergic and inflammatory markers that result in neuronal loss. Neuroinflammation and 5-HT depletion may in turn promote expression of genes involved in tau phosphorylation and aggregation, resulting in tau pathology and the formation of neurofibrillary tangles. We examined mRNA expression of a variety of genes involved in monoaminergic signaling, neuroinflammation and tau phosphorylation and aggregation in the DRN and LC of 4-month-old male htau or C57BL/6J mice (Table 1). In the DRN, we found a significant reduction in Tph2 (t_7_=2.42, p<0.05) and Sert (t_6_=2.69, p<0.05) which is indicative of decreased 5-HT synthesis or 5-HT neurodegeneration. This was accompanied by an increase in TH (t_8_=2.46) which may translate to increased dopamine synthesis (Figure 5A-B). A subset of DRN neurons in the dorsal aspect are dopaminergic and thought to play a role in rebound social behavior after periods of isolation[57]. No significant change in 5-HT receptor expression was observed with the exception of an increase in 5-HT2C (t_6_=5.03, p<0.01) (Figure 5C), which may be in response to reduced 5-HTergic input to GABAergic neurons. These receptors were previously found to regulate GABAergic neurons in the DRN that provide inhibitory input to 5-HT neurons[58], so an increase in this receptor may further reduce 5-HT neuronal activity. Interestingly, there was no change in MAO-A which metabolizes 5-HT to 5-HT1AA (Figure 5B), but there was an increase in indoleamine 2,3 dioxygenase 1 (IDO-1; t_7_=3.03, p<0.05) (Figure 5D), the rate-limiting enzyme in the conversion of tryptophan to kynurenine. Since Tph2 competes with IDO-1 for conversion of Tph2 to 5-HT, an increase in this enzyme together with a decrease in Tph2 would be expected to reduce 5-HT biosynthesis. IDO-1 levels are known to increase in response to inflammatory stimuli[59], and we also found that a number of pro-inflammatory genes were upregulated in htau mice including IL-1a (t_7_=3.42, p<0.05), IL-1b (t_7_=2.62, p<0.05), IL-6 (t_6_=3.60, p<0.05), IL-10R (t_6_=2.49, p<0.05), Cx3CL1 (t_6_=2.59, p<0.05) and Cx3CR1 (t_8_=2.65, p<0.05) (Figure 5E). This increased neuroinflammation may also have deleterious effects on 5-HT neurons by causing excitotoxicity[59]. We also saw changes in genes that modulate tau phosphorylation and aggregation (Figure 5F). There was an increase in fyn kinase (t_6_=4.17, p<0.01) which was previously shown to promote tau phosphorylation[60,61] and in transglutaminase-2 (t_6_=2.92, p<0.05), an enzyme that crosslinks proteins at lysine and glutamine residues and has been implicated in tau aggregation in AD[62–66]. We also observed an upregulation of amyloid precursor protein (APP, t_7_=3.28, p<0.05) and heat shock factor 1 (HSF-1, t_8_=2.49, p<0.05). HSF-1 was recently found to mediate the neuroprotective effects of 5-HT in response to stress[67,68], so upregulation of this gene may be compensatory.

**Figure 5:**
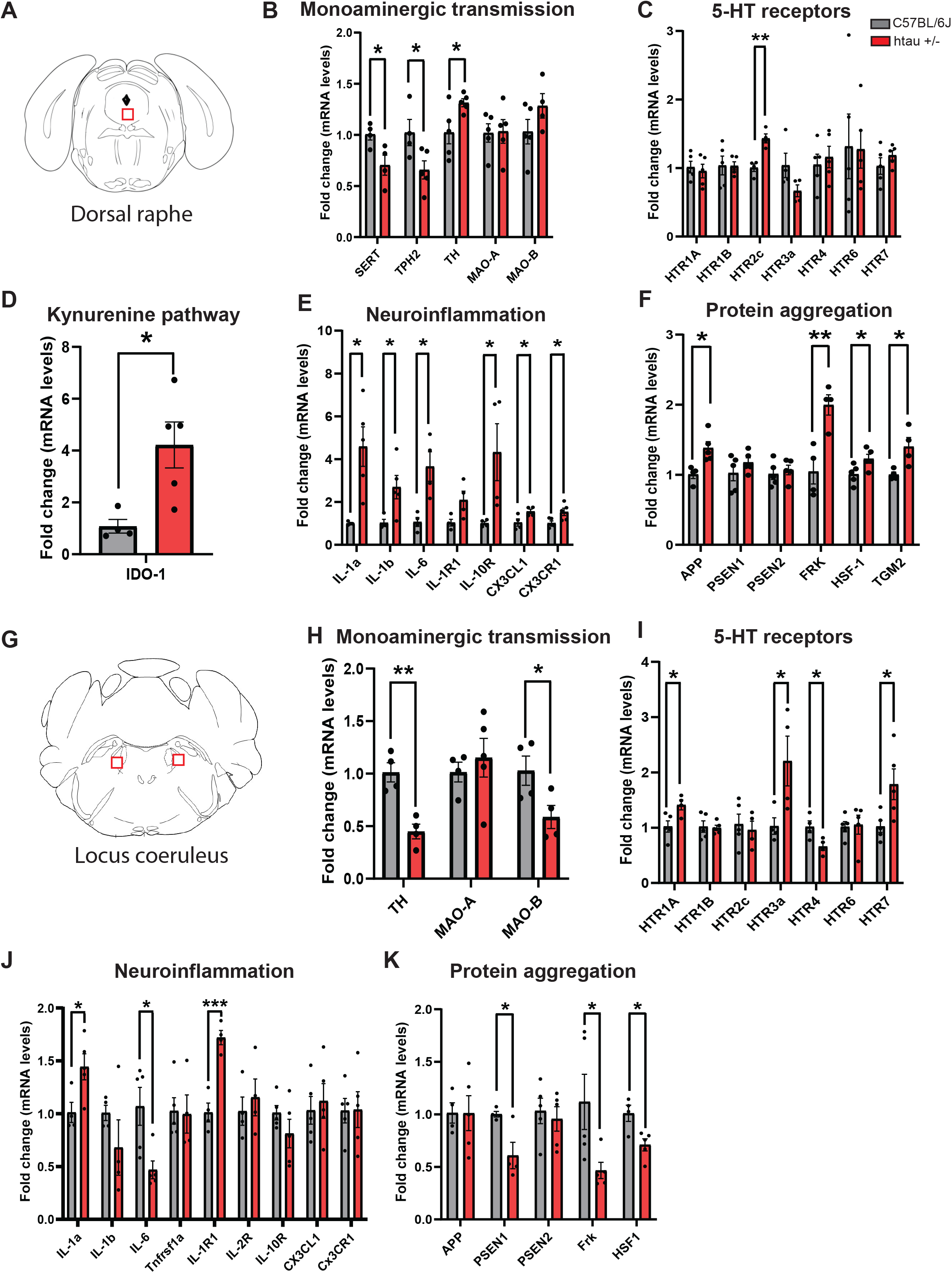
Dysregulation of genes expression in the DRN and LC of htau mice at 4 months. (A) Schematic of tissue extraction in DRN for RT-PCR analysis and results for (B) Monoaminergic genes, (C) 5-HT receptors, (D) IDO-1, (E) Neuroinflammatory genes and (F) Protein aggregation pathways. (G) Schematic of tissue extraction in LC for RT-PCR and results for (H) Monoaminergic transmission (I) 5-HT receptors, (J) Neuroinflammation and (K) Protein aggregation pathways. *p<0.05, **p<0.01, ***p<0.001.

In the LC, there was a significant reduction in tyrosine hydroxylase (TH) expression (t_6_=4.87, p<0.01) and MAO-B (t_6_=2.50, p<0.05) (Figure 5H). This reduced TH gene expression may account for the reduction in TH optical density that we observed in the LC. Interestingly, MAO-A which metabolizes norepinephrine is unchanged, whereas MAO-B metabolizes dopamine and is downregulated. There is also a significant increase in several 5-HT receptors including 5-HT1A (t_7_=2.79, p<0.05), 5-HT3A (t_6_=2.48, p<0.05), and 5-HT7(t_8_=2.55, p<0.05), all of which may be in response to reduced 5-HT input from the DRN (Figure 5I). In the LC, 5-HT3A receptors can stimulate local NE release in the LC which reduces firing of noradrenergic neurons, causing a reduction in distal NE release in the PFC [69]. Loss of noradrenergic neurons or inhibition of their firing in the LC may play a role in the later development of anxiety-like behaviors in htau mice. In contrast, levels of 5-HT4 mRNA in the LC were reduced (t_6_=2.75, p<0.05). 5-HT4 receptors were recently shown to mediate the neuroprotective effects of 5-HT on HSF-1[67], which was also reduced ((t_7_=3.18, p<0.05) and may reflect a loss of 5-HT input to this area. We also observed an increase in pro-inflammatory genes IL-1a (t_7_=2.66, p<0.05) and IL-1R1 (t_6_=6.35, p<0.001), although some cytokine genes like IL-6 were downregulated (t_8_=3.02, p<0.05) (Figure 5J). There was also a decrease in fyn kinase (t_8_=2.38, p<0.05), which is interesting in view of the robust increase in hyperphosphorylated tau in this region (Figure 5K). There was also a decrease in presenilin 1 (PSEN1; t_6_=3.05, p<0.05), which is one of the core proteins in the γ-secretase complex that generates β-amyloid from APP. While we did not directly stain for β-amyloid in this study, it has been reported that amyloid plaques are a rare occurrence in the raphe nuclei of AD patients and likely in the LC as well [21].

### Reduced excitability of 5-HT neurons in the DRN at 4 months

The reduction in 5-HT-expressing neurons in the DRN and optical density may be indicative of neurodegeneration, which can translate to reduced neuronal activity and excitability. We used whole-cell patch clamp electrophysiology to record from 5-HT neurons in the DRN from male 4-month-old C57BL/6J and htau mice (Figure 6A-B). Biocytin was included in the intracellular solution for post-hoc identification of 5-HT neurons, which were analyzed separately. While the resting membrane potential did not differ significantly between groups (Figure 6C), there was a significant decrease in the input resistance (t_34_=2.14, p<0.05) and an increase in the rheobase or action potential threshold (t_34_=2.24, p<0.05) in confirmed 5-HT neurons from htau mice that were initially at their resting membrane potential (RMP; Figure D-F), suggesting that these neurons are less excitable. We also examined the firing frequency during a series of current injections from 0-200 pA starting at RMP and found a main effect of current (F_20,680_=67.41, p<0.001) and a significant interaction between genotype and current (F_20,680_=1.63, p<0.05), suggesting that the frequency-current relationship was significantly modified in the htau mice (e.g. they fire fewer action potentials at higher current steps) (Figure 6G). When the cells were held at −70 mV, we did not observe a significant difference in input resistance but there was a significant increase in the rheobase (t_34_=2.35, p<0.05) and a significant interaction between genotype and current injection (F_20,680_=1.76, p<0.05) as well as a main effect of current (F_20,680_=35.32, p<0.001) (Figure 6H-K). In total, these data support the idea that 5-HT neuronal excitability is reduced in htau mice at 4 months of age.

**Figure 6:**
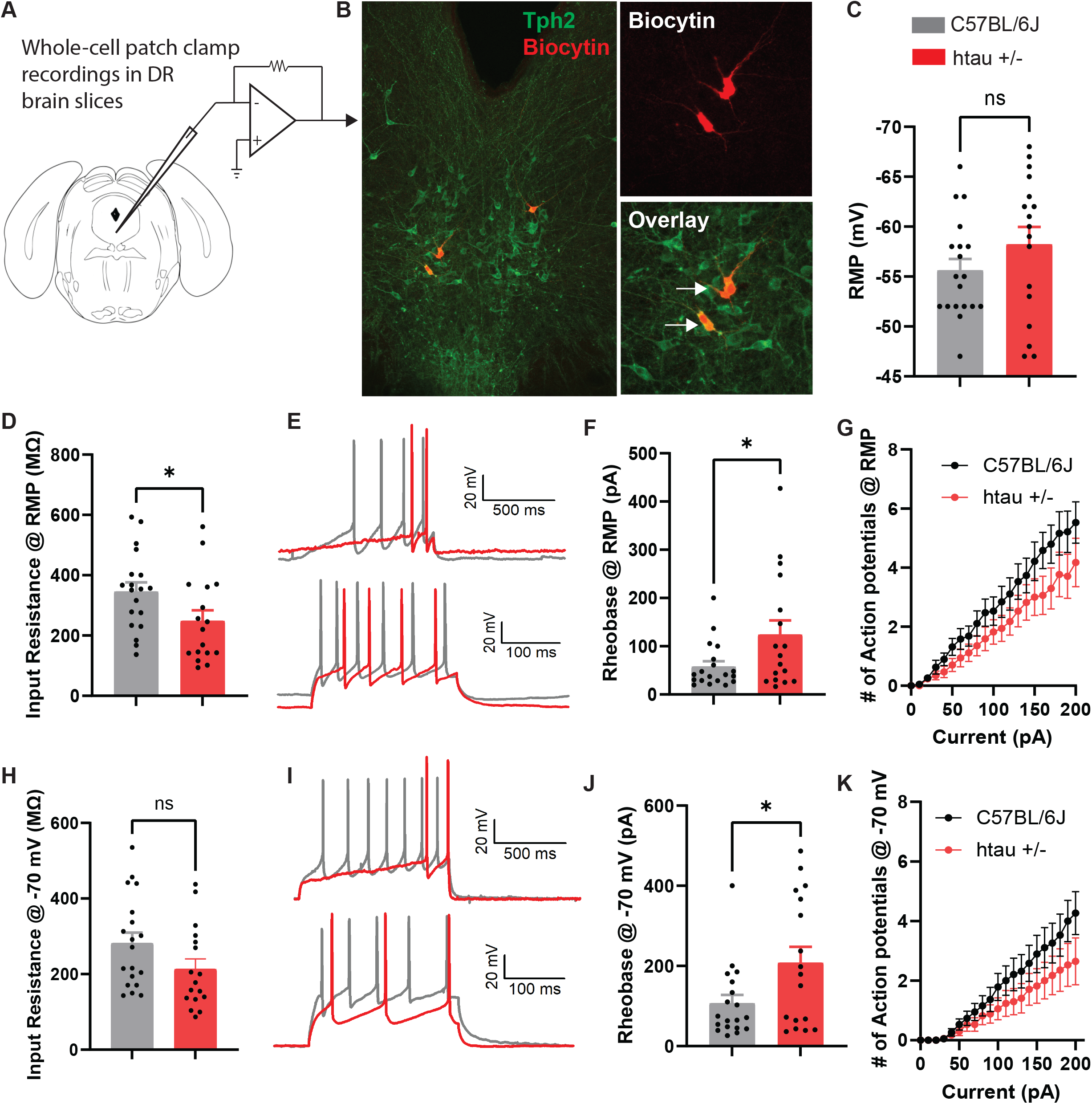
Reduction in 5-HT neuronal excitability in htau mice at 4 months. (A) Schematic of patch clamp recordings in the DRN. (B) Confocal images of biocytin-labeled cells (red) in the DRN overlaid with Tph2 staining (green) indicating that recorded cells were 5-HT neurons. Intracellular recordings in 5-HT neurons from 4-month-old C57 and htau +/- mice with histograms indicating (C) Resting membrane potential (RMP) and (D) Input resistance at RMP. (E) Representative traces of voltage ramps that were used to compute the rheobase and action potential (AP) firing at the 200 pA current step. (F) Rheobase and (G) Current-induced spiking (0-200 pA) from RMP. (H) Input resistance from a holding potential of −70 mV. (I) Representative voltage ramps and AP-current plot at 200 pA from −70 mV. (J) Rheobase and (K) Current induced spiking (0-200 pA) from a holding potential of −70 mV. *p<0.05.

## DISCUSSION

Brainstem monoaminergic nuclei have been implicated in the early stages of AD neuropathology and may be a significant driver of prodromal neuropsychiatric symptoms. Our data in htau mice suggests that tau pathology develops in the DRN at an early age (~ 4 months) and is accompanied by loss of 5-HT neuronal density as well as gene expression changes that reflect a loss of monoaminergic markers, increased inflammatory signaling, and upregulation of genes that promote tau phosphorylation (e.g. fyn kinase and tranglutaminase-2). This coincides with depressive-like behaviors in the social interaction and sucrose preference tests, suggesting that there may be a link between DRN pathology, 5-HT neuronal loss and depression in AD. In addition, there is astrocytic activation in the rostral DRN that may enhance both tau accumulation and 5-HT neuronal loss[59]. Enhanced microglial activity was only observed in the mid and caudal DRN where 5-HT neurons were still intact, suggesting that they may serve a neuroprotective role at this early stage. Interestingly, we did observe hyperactivity in the open field test at 4 months of age, and a previous study suggests that neuroinflammation in the DRN can lead to “manic-like” behavior which includes hyperlocomotion[70]. Hypoexcitability of 5-HT neurons at 4 months also indicates a loss of 5-HT neuronal function which may drive depressive behaviors in these mice. It is also worth noting that depletion of 5-HT neurons occurred at 4 months in our study while others have reported this at 6 months. However, one important distinction is that we stained for 5-HT while this other study probed for Tph, the rate-limiting enzyme in 5-HT biosynthesis[47]. Given that IDO-1 is elevated in the DRN, it is possible that tryptophan is shunted away from the 5-HT biosynthesis and toward the kynurenine pathway. IDO-1 is also upregulated by inflammatory stimuli which are upregulated in the DRN of htau mice at this time point.

In view of accumulating evidence of LC involvement in the early stages of AD, it is surprising that we do not observe a significant reduction in TH-positive neuronal density at the 4-month time point despite the presence of hyperphosphorylated tau in these neurons and a reduction in TH gene expression. This may suggest that TH neuronal degeneration occurs at a later time point than 5-HT neurodegeneration or that 5-HT neurons are more sensitive to the neurotoxic effects of tau aggregation. We also observe anxiety-like behaviors at the 6-month mark which may be the result of neuronal loss or decreased functional output of LC neurons, although we did not explicitly look for these changes at 6 months. It is worth noting that the lack of anxiety-like behavior at 4 months may be attributed to the fact that we used the EPM, which may be less reliable than the EZM that was used at the 6-month mark. In a future study, we would like to test anxiety-like behavior in these mice in the EZM at 4 months of age.

It is also possible that depressive and anxiety-like behaviors in these mice are the result of loss of serotonergic or noradrenergic inputs to other brain regions including the hippocampus and amygdala, which may in turn cause dysregulation of monoaminergic signaling. In a recent study, 5-HT7 receptors in the hippocampus were found necessary for the development of depressive behavior[71], and although we did not explicitly look at the hippocampus, 5-HT7 receptors were upregulated in another region in our study, the LC. Future studies are needed to fully characterize the impact of this early loss of 5-HT neurons in the rostral DRN on downstream 5-HT signaling and receptor expression at neural loci that have been implicated in depression and anxiety.

This study is also the first to establish the htau mouse as a viable model of prodromal AD that could be used to delineate the mechanisms driving early tau accumulation in monoaminergic neurons and their role in the progression of AD. The co-occurrence of depressive behavior, tau pathology and 5-HT dysfunction strongly suggests a role for DRN 5-HT neurons in these early symptoms, although further studies are still needed to establish a causal relationship. This initial investigation lays the foundation for such a study and suggests several targets (e.g. Tph2, Fyn, Tgm2) that could be used to rescue depressive-like behaviors in these mice. There is also indirect evidence to suggest that the LC may be involved in the later development of anxiety-like behaviors at 6 months of age since TH neurons are still intact at 4 months, but reduced TH optical density and increased microglial activity may be a precursor to neurodegeneration at 6 months. Further studies are needed to define changes in TH neuronal density at 6 months of age.

## CONCLUSION

These results support the hypothesis that tau accumulation in the DRN and subsequent loss of serotonergic drive can promote depressive-like behaviors in the prodromal phase of AD prior to the onset of cognitive decline. Further studies in AD mouse models are needed to define a causal relationship between DRN or LC neuropathology and specific AD symptoms.

## DECLARATIONS

### Ethics approval

All procedures on mice in this study were approved by the Institutional Care and Use Committee at the University of Iowa.

### Consent for publication

Not applicable

### Availability of data and materials

All data generated or analyzed during this study are included in this published article and its supplementary information files.

### Authors’ contributions

C.A.M and K.K. designed the studies and wrote the manuscript with editorial help from M.H. K.K also performed behavioral and histological studies and analysis. S.G.P. performed data collection and analysis for RT-PCR, Western blot and histology studies. N.B. performed tissue collection and qRT-PCR analysis of gene expression in the DRN and LC. R.W. performed slice electrophysiology experiments in the DRN, and S.P. performed histological experiments and analysis. M.H. also provided intellectual input into validating the htau line and interpretating the histological results.

## Acknowledgements

We would like to thank Gloria Lee for providing RD3 and RD4 antibodies for Western blot analysis and Gabrielle Bierlein-De La Rosa for technical assistance with behavioral studies.

## REFERENCES

1. Alzheimer’s Association. 2016 Alzheimer’s disease facts and figures. Alzheimers Dement. 2016;12:459–509.

2. Grinberg LT, Rüb U, Ferretti REL, Nitrini R, Farfel JM, Polichiso L, et al. The dorsal raphe nucleus shows phospho-tau neurofibrillary changes before the transentorhinal region in Alzheimer’s disease. A precocious onset? Neuropathol Appl Neurobiol [Internet]. 2009;35:406–16. Available from: http://www.ncbi.nlm.nih.gov/pubmed/19508444

3. Theofilas P, Ehrenberg AJ, Nguy A, Thackrey JM, Dunlop S, Mejia MB, et al. Probing the correlation of neuronal loss, neurofibrillary tangles, and cell death markers across the Alzheimer’s disease Braak stages: a quantitative study in humans. Neurobiol Aging [Internet]. 2018;61:1–12. Available from: http://www.ncbi.nlm.nih.gov/pubmed/29031088

4. Michelsen KA, Prickaerts J, Steinbusch HWM. The dorsal raphe nucleus and serotonin: implications for neuroplasticity linked to major depression and Alzheimer’s disease. Prog Brain Res [Internet]. 2008;172:233–64. Available from: http://www.ncbi.nlm.nih.gov/pubmed/18772036

5. Modrego PJ. Depression in Alzheimer’s Disease. Pathophysiology, Diagnosis, and Treatment. Journal of Alzheimer’s Disease [Internet]. 2010;21:1077–87. Available from: https://www.medra.org/servlet/aliasResolver?alias=iospress&doi=10.3233/JAD-2010-100153

6. Lyketsos CG, Carrillo MC, Ryan JM, Khachaturian AS, Trzepacz P, Amatniek J, et al. Neuropsychiatric symptoms in Alzheimer’s disease. Alzheimer’s & Dementia [Internet]. 2011;7:532–9. Available from: http://doi.wiley.com/10.1016/j.jalz.2011.05.2410

7. Steffens DC, McQuoid DR, Potter GG. Amnestic mild cognitive impairment and incident dementia and Alzheimer’s disease in geriatric depression. Int Psychogeriatr [Internet]. 2014;26:2029–36. Available from: http://www.ncbi.nlm.nih.gov/pubmed/25032667

8. Tao P, Yang S-N, Tung Y-C, Yang M-C. Development of Alzheimer disease in old major depressive patients based upon their health status: A retrospective study in Taiwan. Medicine [Internet]. 2019;98:e15527. Available from: http://www.ncbi.nlm.nih.gov/pubmed/31096454

9. Cohen JY, Amoroso MW, Uchida N. Serotonergic neurons signal reward and punishment on multiple timescales. Elife. 2015;4:e06346.

10. Urban DJ, Zhu H, Marcinkiewcz CA, Michaelides M, Oshibuchi H, Rhea D, et al. Elucidation of the Behavioral Program and Neuronal Network Encoded by Dorsal Raphe Serotonergic Neurons. Neuropsychopharmacology. 2015;

11. Marcinkiewcz CA, Mazzone CM, D’Agostino G, Halladay LR, Hardaway JA, DiBerto JF, et al. Serotonin engages an anxiety and fear-promoting circuit in the extended amygdala. Nature [Internet]. 2016;537:97–101. Available from: http://www.ncbi.nlm.nih.gov/pubmed/27556938

12. Oikonomou G, Altermatt M, Zhang R-W, Coughlin GM, Montz C, Gradinaru V, et al. The Serotonergic Raphe Promote Sleep in Zebrafish and Mice. Neuron [Internet]. 2019;103:686–701.e8. Available from: http://www.ncbi.nlm.nih.gov/pubmed/31248729

13. Paul ED, Lowry C a. Functional topography of serotonergic systems supports the Deakin/Graeff hypothesis of anxiety and affective disorders. J Psychopharmacol [Internet]. 2013;27:1090–106. Available from: http://www.ncbi.nlm.nih.gov/pubmed/23704363

14. Challis C, Beck SG, Berton O. Optogenetic modulation of descending prefrontocortical inputs to the dorsal raphe bidirectionally bias socioaffective choices after social defeat. Front Behav Neurosci. 2014;8:43.

15. Ren J, Friedmann D, Xiong J, Liu CD, Ferguson BR, Weerakkody T, et al. Anatomically Defined and Functionally Distinct Dorsal Raphe Serotonin Sub-systems. Cell [Internet]. 2018; Available from: http://www.ncbi.nlm.nih.gov/pubmed/30146164

16. Weiss JM, Stout JC, Aaron MF, Quan N, Owens MJ, Butler PD, et al. Depression and anxiety: role of the locus coeruleus and corticotropin-releasing factor. Brain Res Bull. 1994;35:561–72.

17. Du X, Yin M, Yuan L, Zhang G, Fan Y, Li Z, et al. Reduction of depression-like behavior in rat model induced by ShRNA targeting norepinephrine transporter in locus coeruleus. Transl Psychiatry. 2020;10:130.

18. Szot P, Franklin A, Miguelez C, Wang Y, Vidaurrazaga I, Ugedo L, et al. Depressive-like behavior observed with a minimal loss of locus coeruleus (LC) neurons following administration of 6-hydroxydopamine is associated with electrophysiological changes and reversed with precursors of norepinephrine. Neuropharmacology. 2016;101:76–86.

19. Bondareff W, Mountjoy CQ, Roth M. Loss of neurons of origin of the adrenergic projection to cerebral cortex (nucleus locus ceruleus) in senile dementia. Neurology. 1982;32:164–8.

20. Zweig RM, Ross CA, Hedreen JC, Steele C, Cardillo JE, Whitehouse PJ, et al. The neuropathology of aminergic nuclei in Alzheimer’s disease. Ann Neurol [Internet]. 1988;24:233–42. Available from: http://www.ncbi.nlm.nih.gov/pubmed/3178178

21. Chen CP, Eastwood SL, Hope T, McDonald B, Francis PT, Esiri MM. Immunocytochemical study of the dorsal and median raphe nuclei in patients with Alzheimer’s disease prospectively assessed for behavioural changes. Neuropathol Appl Neurobiol [Internet]. 2000;26:347–55. Available from: http://www.ncbi.nlm.nih.gov/pubmed/10931368

22. Hendricksen M, Thomas AJ, Ferrier IN, Ince P, O’Brien JT. Neuropathological study of the dorsal raphe nuclei in late-life depression and Alzheimer’s disease with and without depression. Am J Psychiatry [Internet]. 2004;161:1096–102. Available from: http://www.ncbi.nlm.nih.gov/pubmed/15169699

23. Oh J, Eser RA, Ehrenberg AJ, Morales D, Petersen C, Kudlacek J, et al. Profound degeneration of wake-promoting neurons in Alzheimer’s disease. Alzheimers Dement [Internet]. 2019; Available from: http://www.ncbi.nlm.nih.gov/pubmed/31416793

24. Andorfer C, Kress Y, Espinoza M, de Silva R, Tucker KL, Barde Y-A, et al. Hyperphosphorylation and aggregation of tau in mice expressing normal human tau isoforms. J Neurochem [Internet]. 2003;86:582–90. Available from: http://www.ncbi.nlm.nih.gov/pubmed/12859672

25. Polydoro M, Acker CM, Duff K, Castillo PE, Davies P. Age-dependent impairment of cognitive and synaptic function in the htau mouse model of tau pathology. J Neurosci [Internet]. 2009;29:10741–9. Available from: http://www.ncbi.nlm.nih.gov/pubmed/19710325

26. Balasubramanian N, Sagarkar S, Choudhary AG, Kokare DM, Sakharkar AJ. Epigenetic Blockade of Hippocampal SOD2 Via DNMT3b-Mediated DNA Methylation: Implications in Mild Traumatic Brain Injury-Induced Persistent Oxidative Damage. Molecular Neurobiology. 2021;58:1162–84.

27. Andorfer C, Kress Y, Espinoza M, de Silva R, Tucker KL, Barde YA, et al. Hyperphosphorylation and aggregation of tau in mice expressing normal human tau isoforms. J Neurochem. 2003/07/16. 2003;86:582–90.

28. Duff K, Knight H, Refolo LM, Sanders S, Yu X, Picciano M, et al. Characterization of pathology in transgenic mice over-expressing human genomic and cDNA tau transgenes. Neurobiol Dis. 2000/04/28. 2000;7:87–98.

29. Schmittgen TD, Livak KJ. Analyzing real-time PCR data by the comparative CT method. Nature Protocols. 2008;3:1101–8.

30. Li L, Shi R, Gu J, Tung YC, Zhou Y, Zhou D, et al. Alzheimer’s disease brain contains tau fractions with differential prion-like activities. Acta Neuropathol Commun. 2021;9:28.

31. Julien C, Bretteville A, Planel E. Biochemical isolation of insoluble tau in transgenic mouse models of tauopathies. Methods Mol Biol. 2012;849:473–91.

32. Yin X, Jin N, Shi J, Zhang Y, Wu Y, Gong C-X, et al. Dyrk1A overexpression leads to increase of 3R-tau expression and cognitive deficits in Ts65Dn Down syndrome mice. Sci Rep. 2017;7:619.

33. Congdon EE, Lin Y, Rajamohamedsait HB, Shamir DB, Krishnaswamy S, Rajamohamedsait WJ, et al. Affinity of Tau antibodies for solubilized pathological Tau species but not their immunogen or insoluble Tau aggregates predicts in vivo and ex vivo efficacy. Mol Neurodegener. 2016;11:62.

34. Andorfer CA, Davies P. PKA phosphorylations on tau: developmental studies in the mouse. Dev Neurosci. 2000;22:303–9.

35. Shoji H, Miyakawa T. Effects of test experience, closed-arm wall color, and illumination level on behavior and plasma corticosterone response in an elevated plus maze in male C57BL/6J mice: a challenge against conventional interpretation of the test. Mol Brain. 2021;14:34.

36. Coutellier L, Beraki S, Ardestani PM, Saw NL, Shamloo M. Npas4: A Neuronal Transcription Factor with a Key Role in Social and Cognitive Functions Relevant to Developmental Disorders. PLoS ONE. 2012;7.

37. Shepherd JK, Grewal SS, Fletcher A, Bill DJ, Dourish CT. Behavioural and pharmacological characterisation of the elevated “zero-maze” as an animal model of anxiety. Psychopharmacology (Berl). 1994;116:56–64.

38. Gonçalves RA, Wijesekara N, Fraser PE, De Felice FG. Behavioral Abnormalities in Knockout and Humanized Tau Mice. Frontiers in Endocrinology. 2020;11:1–13.

39. Kaidanovich-Beilin O, Lipina T, Vukobradovic I, Roder J, Woodgett JR. Assessment of social interaction behaviors. Journal of Visualized Experiments. 2010;0:1–6.

40. Yang M, Silverman JL, Crawley JN. Automated three-chambered social approach task for mice. Current Protocols in Neuroscience. 2011;

41. Liu MY, Yin CY, Zhu LJ, Zhu XH, Xu C, Luo CX, et al. Sucrose preference test for measurement of stress-induced anhedonia in mice. Nature Protocols. 2018;13:1686–98.

42. Romano A, Pace L, Tempesta B, Lavecchia AM, Macheda T, Bedse G, et al. Depressive-like behavior is paired to monoaminergic alteration in a murine model of Alzheimer’s disease. International Journal of Neuropsychopharmacology. 2014;18:1–12.

43. Flinn JM, Lorenzo Bozzelli P, Adlard PA, Railey AM. Spatial memory deficits in a mouse model of late-onset Alzheimer’s disease are caused by zinc supplementation and correlate with amyloid-beta levels. Frontiers in Aging Neuroscience. 2014;6:1–10.

44. Lippi SLP, Smith ML, Flinn JM. A Novel hAPP/htau Mouse Model of Alzheimer’s Disease: Inclusion of APP With Tau Exacerbates Behavioral Deficits and Zinc Administration Heightens Tangle Pathology. Frontiers in Aging Neuroscience. 2018;10.

45. Gonçalves RA, Wijesekara N, Fraser PE, De Felice FG. Behavioral Abnormalities in Knockout and Humanized Tau Mice. Frontiers in Endocrinology. 2020;11:1–13.

46. Attar A, Liu T, Chan W-TC, Hayes J, Nejad M, Lei K, et al. A Shortened Barnes Maze Protocol Reveals Memory Deficits at 4-Months of Age in the Triple-Transgenic Mouse Model of Alzheimer’s Disease. Coulson EJ, editor. PLoS ONE. 2013;8:e80355.

47. Dengler-Crish CM, Smith MA, Wilson GN. Early Evidence of Low Bone Density and Decreased Serotonergic Synthesis in the Dorsal Raphe of a Tauopathy Model of Alzheimer’s Disease. J Alzheimers Dis [Internet]. 2017;55:1605–19. Available from: http://www.ncbi.nlm.nih.gov/pubmed/27814296

48. Graeff FG, Guimarães FS, de Andrade TG, Deakin JF. Role of 5-HT in stress, anxiety, and depression. Pharmacol Biochem Behav [Internet]. 1996 [cited 2014 Dec 21];54:129–41. Available from: http://www.ncbi.nlm.nih.gov/pubmed/8728550

49. Ren J, Isakova A, Friedmann D, Zeng J, Grutzner SM, Pun A, et al. Single-cell transcriptomes and whole-brain projections of serotonin neurons in the mouse dorsal and median raphe nuclei. Elife [Internet]. 2019;8. Available from: https://elifesciences.org/articles/49424

50. in t’ Veld BA, Ruitenberg A, Hofman A, Launer LJ, van Duijn CM, Stijnen T, et al. Nonsteroidal antiinflammatory drugs and the risk of Alzheimer’s disease. N Engl J Med. 2001;345:1515–21.

51. Heppner FL, Ransohoff RM, Becher B. Immune attack: the role of inflammation in Alzheimer disease. Nat Rev Neurosci. 2015;16:358–72.

52. Condello C, Yuan P, Schain A, Grutzendler J. Microglia constitute a barrier that prevents neurotoxic protofibrillar Aβ42 hotspots around plaques. Nat Commun. 2015;6:6176.

53. Hansen D v, Hanson JE, Sheng M. Microglia in Alzheimer’s disease. J Cell Biol. 2018;217:459–72.

54. Carrero I, Gonzalo MR, Martin B, Sanz-Anquela JM, Arévalo-Serrano J, Gonzalo-Ruiz A. Oligomers of β-amyloid protein (Aβ1-42) induce the activation of cyclooxygenase-2 in astrocytes via an interaction with interleukin-1β, tumour necrosis factor-α, and a nuclear factor κ-B mechanism in the rat brain. Exp Neurol. 2012;236:215–27.

55. Navarro V, Sanchez-Mejias E, Jimenez S, Muñoz-Castro C, Sanchez-Varo R, Davila JC, et al. Microglia in Alzheimer’s Disease: Activated, Dysfunctional or Degenerative. Front Aging Neurosci. 2018;10:140.

56. Heneka MT, Nadrigny F, Regen T, Martinez-Hernandez A, Dumitrescu-Ozimek L, Terwel D, et al. Locus ceruleus controls Alzheimer’s disease pathology by modulating microglial functions through norepinephrine. Proc Natl Acad Sci U S A. 2010;107:6058–63.

57. Matthews GA, Nieh EH, vander Weele CM, Halbert SA, Pradhan R v, Yosafat AS, et al. Dorsal Raphe Dopamine Neurons Represent the Experience of Social Isolation. Cell. 2016;164:617–31.

58. Spoida K, Masseck O a, Deneris ES, Herlitze S. Gq/5-HT2c receptor signals activate a local GABAergic inhibitory feedback circuit to modulate serotonergic firing and anxiety in mice. Proc Natl Acad Sci U S A [Internet]. 2014 [cited 2014 Nov 24];111:6479–84. Available from: http://www.pubmedcentral.nih.gov/articlerender.fcgi?artid=4035925&tool=pmcentrez&rendertype=abstract

59. Hochstrasser T, Ullrich C, Sperner-Unterweger B, Humpel C. Inflammatory stimuli reduce survival of serotonergic neurons and induce neuronal expression of indoleamine 2,3-dioxygenase in rat dorsal raphe nucleus organotypic brain slices. Neuroscience [Internet]. 2011;184:128–38. Available from: http://www.ncbi.nlm.nih.gov/pubmed/21501664

60. Lee G, Thangavel R, Sharma VM, Litersky JM, Bhaskar K, Fang SM, et al. Phosphorylation of tau by fyn: implications for Alzheimer’s disease. J Neurosci. 2004;24:2304–12.

61. Liu G, Fiock KL, Levites Y, Golde TE, Hefti MM, Lee G. Fyn depletion ameliorates tauP301L-induced neuropathology. Acta Neuropathol Commun. 2020;8:108.

62. Norlund MA, M. Lee J, Zainelli GM, Muma NA. Elevated transglutaminase-induced bonds in PHF tau in Alzheimer’s disease. Brain Research. 1999;851:154–63.

63. Zemaitaitis MO, Lee JM, Troncoso JC, Muma NA. Transglutaminase-Induced Cross-Linking of Tau Proteins in Progressive Supranuclear Palsy. Journal of Neuropathology & Experimental Neurology. 2000;59:983–9.

64. Halverson RA, Lewis J, Frausto S, Hutton M, Muma NA. Tau protein is cross-linked by transglutaminase in P301L tau transgenic mice. J Neurosci. 2005;25:1226–33.

65. Grierson AJ, Johnson G v, Miller CC. Three different human tau isoforms and rat neurofilament light, middle and heavy chain proteins are cellular substrates for transglutaminase. Neurosci Lett [Internet]. 2001;298:9–12. Available from: http://www.ncbi.nlm.nih.gov/pubmed/11154823

66. Tucholski J, Kuret J, Johnson G v. Tau is modified by tissue transglutaminase in situ: possible functional and metabolic effects of polyamination. J Neurochem. 1999;73:1871–80.

67. Das S, Ooi FK, Cruz Corchado J, Fuller LC, Weiner JA, Prahlad V. Serotonin signaling by maternal neurons upon stress ensures progeny survival. Elife. 2020;9.

68. Cruz-Corchado J, Ooi FK, Das S, Prahlad V. Global Transcriptome Changes That Accompany Alterations in Serotonin Levels in Caenorhabditis elegans. G3 (Bethesda). 2020;10.

69. Ortega JE, Mendiguren A, Pineda J, Meana JJ. Regulation of central noradrenergic activity by 5-HT(3) receptors located in the locus coeruleus of the rat. Neuropharmacology. 2012;62:2472–9.

70. Howerton AR, Roland A v, Bale TL. Dorsal raphe neuroinflammation promotes dramatic behavioral stress dysregulation. J Neurosci. 2014;34:7113–23.

71. Bijata M, Bączyńska E, Müller FE, Bijata K, Masternak J, Krzystyniak A, et al. Activation of the 5-HT7 receptor and MMP-9 signaling module in the hippocampal CA1 region is necessary for the development of depressive-like behavior. Cell Rep. 2022;38:110532.

