## Supplementary material for "Behavioral dysregulation and monoaminergic deficits precede memory impairments in human tau-overexpressing (htau) mice": Tables

**Table 1: Primers used for quantitating mRNA expression using RT-qPCR**

| Gene Name | Forward/Reverse (5’-3’) | Sequence |
| --- | --- | --- |
| *β-actin* | Forward  Reverse | CCAGCCTTCCTTCTTGGGTA  GAGGTCTTTACGGATGTCAACG |
| *Sert* | Forward  Reverse | CAAAACGTCTGGCAAGGTGG  ACACCCCTGTCTCCAAGAGT |
| *Tph2* | Forward  Reverse | GACCCAAAGACGACCTGCTT  CTGCGTGTAGGGGTTGAAGT |
| *Th* | Forward  Reverse | TACTTTGTGCGCTTCGAGGT  GGAACCTTGTCCTCTCTGGC |
| *Mao-a* | Forward  Reverse | GTATGTGAGGCAGTGTGGAGG  CCCCAAGGAGGACCATTATCTG |
| *Mao-b* | Forward  Reverse | ATTCCACCTGCTTTGGGCAT  TGAACCCAAAGGCACACGA |
| *Htr1a* | Forward  Reverse | TACTCCACTTTCGGCGCTTT  GGCTGACCATTCAGGCTCTT |
| *Htr1b* | Forward  Reverse | ACCCTTCTTCTGGCGTCAAG  ACCGTGGAGTAGACCGTGT |
| *Htr2c* | Forward  Reverse | GTGCCCGTTTTTCATCACCAA  AGGAGGCTTTTTGTCTGGCTT |
| *Htr3a* | Forward  Reverse | CCATCTTCATTGTGCGGCTG  CTTGTTGGCTTGGAAGGTGG |
| *Htr4* | Forward  Reverse | GGAGTGTGCCAGGAGATCAG  AACCACTGCAAGGAACGTGA |
| *Htr6* | Forward  Reverse | AGTGGGAGGTGGTAGGTCTC  GGGCTGAGGACTGATTGCTT |
| *Htr7* | Forward  Reverse | AAGTTCTCAGGCTTCCCACG  TTCGCACACTCTTCCACCTC |
| *Il-1a* | Forward  Reverse | TTGCTGAAGGAGTTGCCAGA  GCACCCGACTTTGTTCTTTGG |
| *Il-1b* | Forward  Reverse | GCCACCTTTTGACAGTGATGAG  AAGGTCCACGGGAAAGACAC |
| *Il-6* | Forward  Reverse | GAGACTTCCATCCAGTTGCCT  TCCTCTGTGAAGTCTCCTCTCC |
| *Tnfrsf1a* | Forward  Reverse | AGCCACACCCACAACCTTAG  CCCCTTAGAGACCTTTGCCC |
| *Il-1r1* | Forward  Reverse | ACTTGAGGAGGCAGTTTTCGT  GTCAATCTCCAGCGACAGCA |
| *Il-2r* | Forward  Reverse | TGAAGTGTGGGAAAACGGGG  GCAGGAAGTCTCACTCTCGG |
| *Il-10r* | Forward  Reverse | GCGTGACTCTGAAAGCAATGG  GCAGCACCTTGACACAAAACT |
| *Cx3cl1* | Forward  Reverse | GAGAGTGAGGAAGCCAACCC  AAAGTCCGATGACGGGTGTC |
| *Cx3cr1* | Forward  Reverse | GGGTTTGGTGAGTCCTGGTT  CAAGGAATGGACACCCGACA |
| *App* | Forward  Reverse | TTCGCTGACGGAAACCAAGA  TTTCGGTATTGGCTGGCACA |
| *Psen1* | Forward  Reverse | CTCCTGCTCGCCATTTTCAAG  CACAAGGTAATCCGTGGCGA |
| *Psen2* | Forward  Reverse | ACACTGAGAAGAACGGGCAG  AGGAGCATCAGGGAGGACAT |
| *Frk* | Forward  Reverse | AGCAGGTCAGGAAGAAGCAC  CTCACCATACCTCCCGCTTC |
| *Hsf1* | Forward  Reverse | AGAGGAAAGTGACCAGCGTG  ACAACTTTTTGCTGCTGGGC |

| **Stain** | **Antibody** | **Concentration** |
| --- | --- | --- |
| 5HT-AT8 | Goat anti-5HT  (Immunostar; catalog # 20079) | 1:2000 |
|  | Mouse anti-AT8  (Thermofisher, catalog # MN1020) | 1:200 |
|  | 405 Donkey anti-Goat  (Jackson ImmunoResearch; catalog # 715-475-003) | 1:500 |
|  | Cy3 Donkey anti-Mouse  (Jackson ImmunoResearch; catalog # 711-165-151) | 1:500 |
| TH-AT8 | Rabbit anti-TH (Novus; catalog # NB300-109) | 1:1000 |
|  | Mouse anti-AT8  (Thermofisher, catalog # MN1020) | 1:200 |
|  | 405 Donkey anti-Rabbit  (Abcam; catalog # ab175651) | 1:500 |
|  | Cy3 Donkey anti-Mouse  (Jackson ImmunoResearch; catalog # 715-165-151) | 1:500 |
| 5HT-iba1-GFAP | Goat anti-5HT (Immunostar; catalog # 20079) | 1:2000 |
|  | Rabbit anti-iba1  (Abcam; catalog # ab178846) | 1:200 |
|  | Chicken anti-GFAP  (Abcam; catalog # ab4674) | 1:6000 |
|  | 405 Donkey anti-Goat  (Jackson ImmunoResearch; catalog # 715-475-003) | 1:500 |
|  | Cy3 Donkey anti-Rabbit  (Jackson ImmunoResearch; catalog # 711-165-152) | 1:500 |
|  | 647 Donkey anti-Chicken  (Jackson ImmunoResearch; catalog # 703-605-155) | 1:500 |
| TH-iba1-GFAP | Mouse anti-TH  (Millipore Sigma; catalog # MAB318) | 1:200 |
|  | Rabbit anti-iba1  (Abcam; catalog # ab178846) | 1:200 |
|  | Chicken anti-GFAP  (Abcam; catalog # ab4674) | 1:6000 |
|  | 405 Donkey anti-Mouse  (Invitrogen; catalog # A48257) | 1:500 |
|  | Cy3 Donkey anti-Rabbit  (Jackson ImmunoResearch; catalog # 711-165-152) | 1:500 |
|  | 647 Donkey anti-Chicken  (Jackson ImmunoResearch; catalog # 703-605-155) | 1:500 |

**Table 2: Antibodies used in immunohistochemistry**
